# Anti-PD-1 treatment as a neurotherapy to enhance neuronal excitability, synaptic plasticity and memory

**DOI:** 10.1101/870600

**Authors:** Junli Zhao, Ru-Rong Ji

## Abstract

Emerging immunotherapy using anti-PD-1 antibodies have improved survival in cancer patients by immune activation. Here we show that functional PD-1 receptor is present in hippocampal CA1 neurons and loss of PD-1 or local anti-PD-1 treatment with nivolumab enhances neuronal excitability and long-term potentiation in CA1 neurons, leading to enhanced learning and memory. Our findings suggest that anti-PD-1 antibody also acts as a neurotherapy for improving memory and counteracting cognitive decline.

---

Programmed cell death protein 1 (PD-1) is an immune checkpoint inhibitor expressed by immune cells such as T cells and serves as a primary target of immunotherapy agents that fight cancer ^1, 2^. New evidence suggests that anti-PD-1 also has an active role in the nervous system. Schwartz and colleagues demonstrated that systemic PD-1 blockade promoted clearance of cerebral amyloid-β plaques and improve memory in mouse models of Alzheimer's disease (AD)^3, 4^ through brain recruitment of peripheral macrophages in the central nervous system (CNS). However, Latta-Mahieu et al. reported that systemic anti-PD-1 treatment failed to affect Aβ pathology in 3 different models of AD^5^. Obst et al. demonstrated that PD-1 deficiency is not sufficient to induce myeloid mobilization to the brain during chronic neurodegeneration^6^. These studies suggested that anti-PD-1 treatment may improve memory through different mechanisms. Recently, we demonstrated for the first time that functional PD-1 is present in neurons^7^. Our previous study showed that PD-1 is expressed by primary sensory neurons of the peripheral nervous system (PNS) and activation of PD-1 on sensory neurons by its ligand PD-L1 suppressed nociceptor excitability and nociceptive behaviors in mice^7^. In this study, we tested the hypothesis that functional PD-1 is also present in CNS neurons. Furthermore, we postulated that *Pd1* deficiency or anti-PD-1 treatment would regulate learning and memory via neuromodulation, i.e. “neurotherapy”.

As a first step to address the role of PD-1 in learning and memory, we compared new object learning in wild-type (WT) and knockout mice lacking *Pdcd1* (*Pd1*^−/−^) of young adults (2-3 months, **Supplementary Fig. 1a**). Notably, *Pd1* gene is partially deleted in *Pd1* ^−/−^ mice: 2 of 5 exons (exon-2 and exon-3) were deleted, leading to a functional loss of PD-1^8^. Compared with WT mice, *Pd1*^−/−^ mice exhibited increased discrimination index at two different time points (0.5 and 24 h, *P*<0.05, **Fig. 1a**), indicating increased capability of new object learning. Water maze test further revealed that the latency to locate the platform was shorter in KO mice (*P*<0.001, **Fig. 1b**, **Supplementary Fig. 1b**) and the number of crossing was enhanced in KO mice (*P*<0.01, vs. WT, **Fig. 1c**). Thus, loss of PD-1 can enhance learning and memory in naïve mice without neurodegeneration.

**Fig.1.**
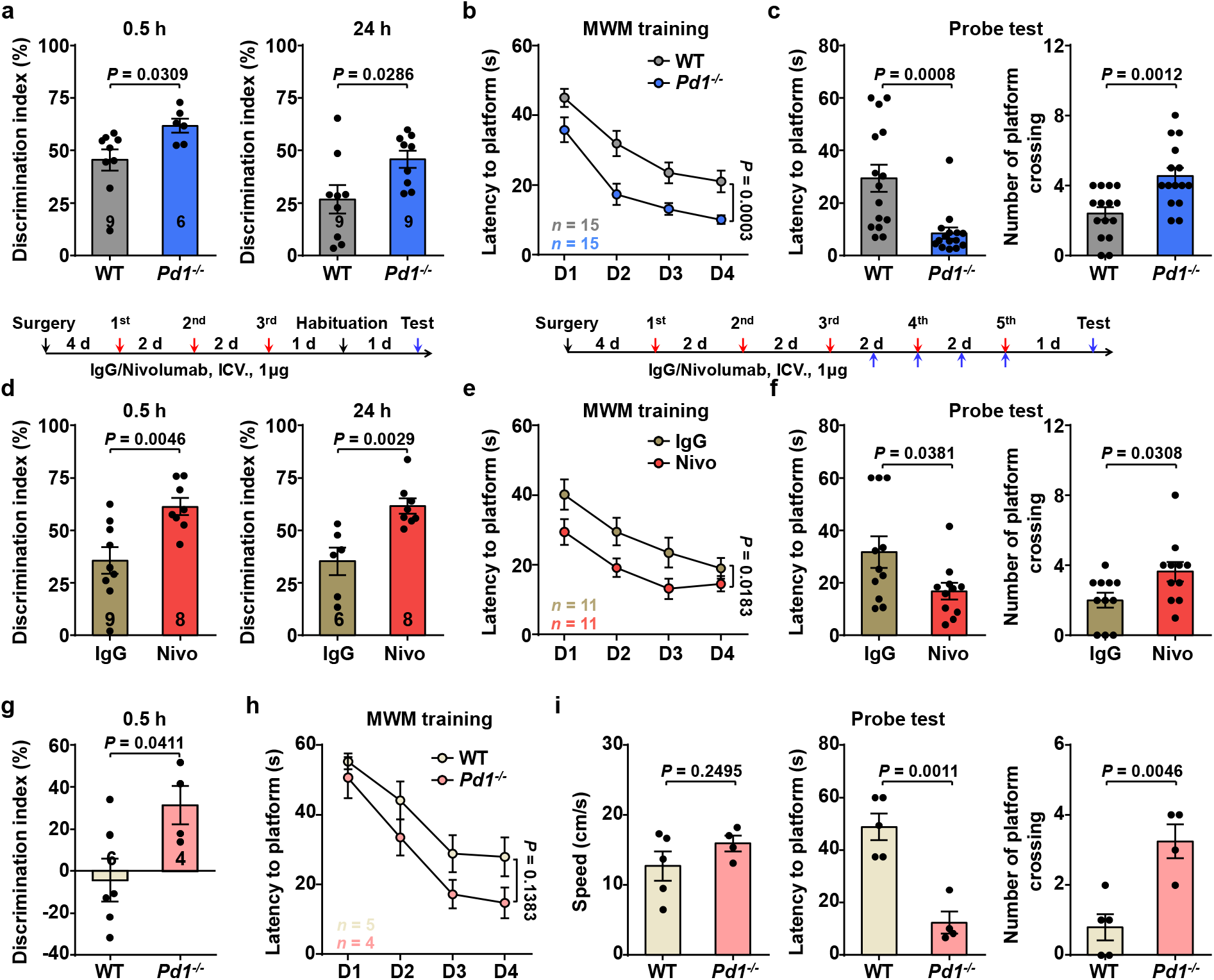
Enhanced learning and memory in *Pd1*^−/−^ mice and wild-type mice treated with intraventricular nivolumab. **(a)** Novel object recognition test at 0.5 h and 24 h in WT and *Pd1*^−/−^ mice. **(b, c)** *Pd1*^−/−^ mice exhibited enhanced learning and memory in Morris water maze test. **(b)** During training phase, both WT and *Pd1*^−/−^ mice showed reduced latency to locate the platform, but *Pd1*^−/−^ mice exhibited a greater improvement. **(c)** In probe test on day 5, *Pd1*^−/−^ mice exhibited shorter latency to locate the platform and more platform zone crossing. **(d)** Novel object recognition test after ICV infusion of nivolumab and control IgG (3 × 1 μg, given every other day) in WT mice. (**e, f**) Morris water maze test after ICV infusion of nivolumab or control IgG (5 × 1 μg, every other day) in WT mice. **(e)** During training phase, both IgG- and nivolumab decreased latency to locate the platform, but nivolumab-treated mice exhibited a greater improvement. **(f)** In probe test on day 5, nivolumab-treated mice decreased latency to locate the platform and increased platform zone crossing. **(g)** 10-month od WT mice showed declined learning and memory ability in novel object recognition test, but 10-month old *Pd1*^−/−^ mice kept the learning ability. **(h, i)** Morris water maze test shows that 10-month old *Pd1*^−/−^ mice have almost intact learning and memory. **(h)** During training phase, both WT and *Pd1*^−/−^ mice showed improved latency to locate the platform. **(i)** In probe test on day 5, WT and KO mice showed no significant difference in swimming speed (left), but *Pd1*^−/−^ mice exhibited a shorter latency to locate the platform (middle) and more platform zone crossing (right). Two-tailed unpaired t test for **a, c, d, f, g** and **i**. Two-way ANOVA with Bonferroni post hoc test for **b, e, h**. Data were expressed as mean ± s.e.m.

To circumvent developmental compensation in *Pd1*^−/−^ mice, we examined whether central PD-1 blockade in WT mice could recapitulate the phenotypes observed in KO mice. Nivolumab is a FDA-approved fully humanized IgG4 monoclonal antibody and binds mouse PD-1 to produce functional blockade in behavioral assays in mice^7^. Intraventricular (ICV) administration of nivolumab at a very low dose (1 μg ≈ 7 pmol, 3 × 1 μg, every other day) significantly enhanced new object learning, as compared to human IgG4 control (*P*<0.01, **Fig. 1d**). ICV administration of nivolumab (5 × 1 μg, every other day) also enhanced memory in the water maze test (*P*<0.05, vs. control IgG4, **Fig. 1e,f**). Notably, swimming speed was slightly increased after *Pd1* deletion but not after nivolumab treatment (**Supplementary Fig. 1c, d**), suggesting that memory improvement is not a result of enhanced swimming capability.

Next, we investigated learning and memory in 10 month-old WT and KO mice. Compared with young adult mice (2-3 months), 10 month-old WT mice showed decreased discrimination index (45% vs. −4%, *P*<0.05, **Fig. 1d, g, Supplementary Fig. 1e**), indicating age-related cognitive decline in mice. However, 10 month-old KO mice had significant higher learning index (*P*<0.05 vs. WT old mice, **Fig. 1g**). 10 month-old WT mice also showed declined memory capacity in water maze test (*P*=0.0546, vs. young mice, **Supplementary Fig. 1f**). Strikingly, 10 month-old-KO mice were protected from memory deterioration, showing significantly shorter latency to locate the platform (*P*<0.01 vs. old-WT mice, **Fig. 1h, i**). Memory was almost intact in 10 month-old-KO mice when compared to young-KO mice (latency: 12 s in 10 month-old-KO mice vs. 9s in young-KO mice, **Fig. 1c, i**, and **Supplementary Fig. 1f**). Notably, the swimming speed between 10 month-old WT and KO mice was comparable (**Fig. 1i**, left). Taken together, these data suggest that PD-1 is an endogenous inhibitor of learning and memory in both young and 10 month-old mice.

Hippocampal CA1 neurons play critical roles in learning and memory ^9, 10^. To define neuronal mechanisms by which PD-1 regulates learning and memory, we examined neuronal excitability in CA1 neurons using brain slice preparation. Patch-clamp recordings showed that *Pd1*-deficient neurons had increased resting membrane potential (RMP, *P*<0.05, WT vs. KO, **Fig. 2a**). *Pd1*-deficient neurons showed no changes in AP amplitude and duration (**Supplementary Fig. 2a**) but fired more action potentials (AP) following a ramp of 0-130 pA current injection (*P*<0.0001, WT vs. KO, **Fig. 2b, Supplementary Fig. 2b**). Intriguingly, spontaneous discharges was found in 40% of *Pd1*-deficient CA1 neurons (8/20) but none were observed in 20 WT neurons (**Fig. 2c**). Thus, loss of PD-1 results in an increase in neuronal excitability in CA1 neurons. Strikingly, this increase was also found in WT mice following anti-PD-1 treatment. Acute treatment of brain slices with nivolumab (7 nM, 2 h) did not change RMP but significantly increased AP number (*P*<0.01, **Fig. 2d, e**) and AP frequency (*P*<0.05, **Supplementary Fig. 2c**) in CA1 neurons. These findings suggest a cell-autonomous PD-1 signaling mechanism within hippocampal neurons.

**Fig. 2:**
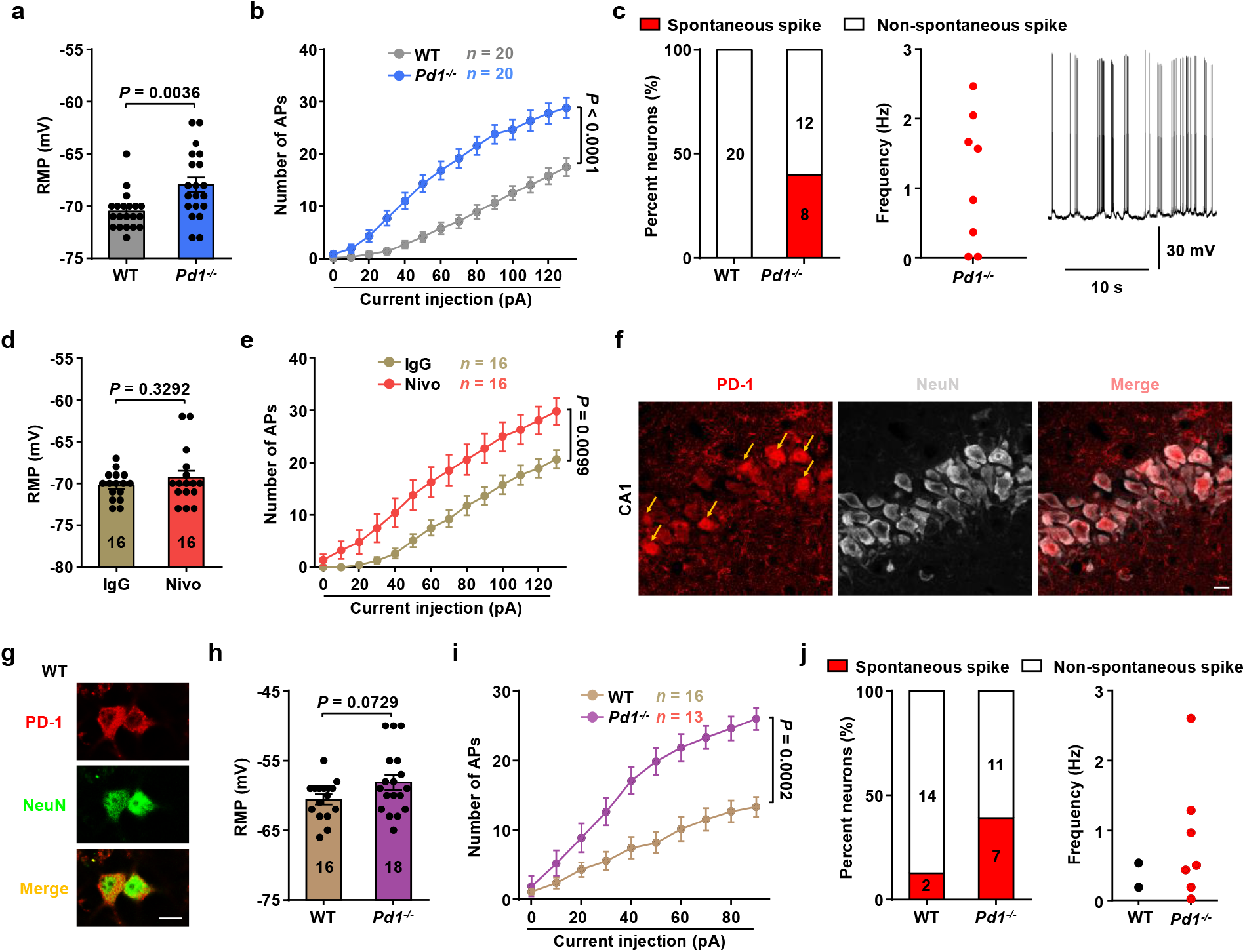
Enhanced excitability in hippocampal CA1 neurons with *Pd1*^−/−^ deficiency or after PD-1 blockade with nivolumab. **(a, b)** Altered RMP and increased excitability in CA1 neurons in brain slices of *Pd1*^−/−^ mice. **(a)** RMP in WT and *Pd1*^−/−^ mice. **(b)** Number of action potentials evoked by current injection in WT and *Pd1*^−/−^ mice. **(c)** Spontaneous discharges in *Pd1*-deficient CA1 neurons of brain slices. Note that 40% (8/20, left) CA1 neurons showed spontaneous spikes at a frequency of 0.02-2.5 Hz. **(d)** RMP in WT-CA1 neurons after treatment of IgG and nivolumab (7 nM, 2 h). **(e)** Number of action potentials evoked by current injection in WT neurons after treatment of IgG- and nivolumab (7 nM, 2 h) in brain slices. **(f)** Immunohistochemistry showing PD-1 expression in CA1 neurons of WT mice. Arrows indicate PD-1^+^ neurons. Scale bar, 10 μm. **(g-j)** Histochemical and electrophysiological characterization of hippocampal neurons in primary cultures (DIV 7-8) prepared from E17-19 mice. **(g)** Immunocytochemistry showing PD-1 and NeuN double staining. Scale bar, 10 μm. **(h)** RMP from WT and *Pd1*^−/−^ neurons. **(i)** Number of action potentials evoked by current injection in WT and *Pd1*^−/−^ hippocampal neurons. **(j)** Spontaneous spikes in primary hippocampal neurons of WT and KO mice. Note that 12.5% (2/16) of WT neurons and 38.9% (7/18) of *Pd1*^−/−^ neurons showed spontaneous spike (0.01-2.6 Hz), respectively. Two-tailed unpaired t test for **a, d, h.** Two-way ANOVA with Bonferroni post hoc test for **b, e, i**. Data were expressed as mean ± s.e.m.

We then examined PD-1 expression and found that hippocampal neurons express PD-1 protein or *Pd1* mRNA. Double staining revealed that PD-1 is expressed by CA1 neurons that co-express the neuronal marker NeuN (**Fig. 2f**). The PD-1 staining was abolished by the blocking peptide^7^ and lost in *Pd1*-KO mice (**Supplementary Fig. 3b,c**). PD-1 was expressed by 20-40% CA1 and CA3 neurons (**Extended Data Fig. 3a, d**). Consistently, in situ hybridization revealed *Pd1* mRNA expression in CA1 and CA3 (**Supplementary Fig. 3e**). Immunocytochemistry in cultured hippocampal neurons also showed PD-1 expression in cytoplasm/surface (**Fig. 2g**).

**Fig. 3:**
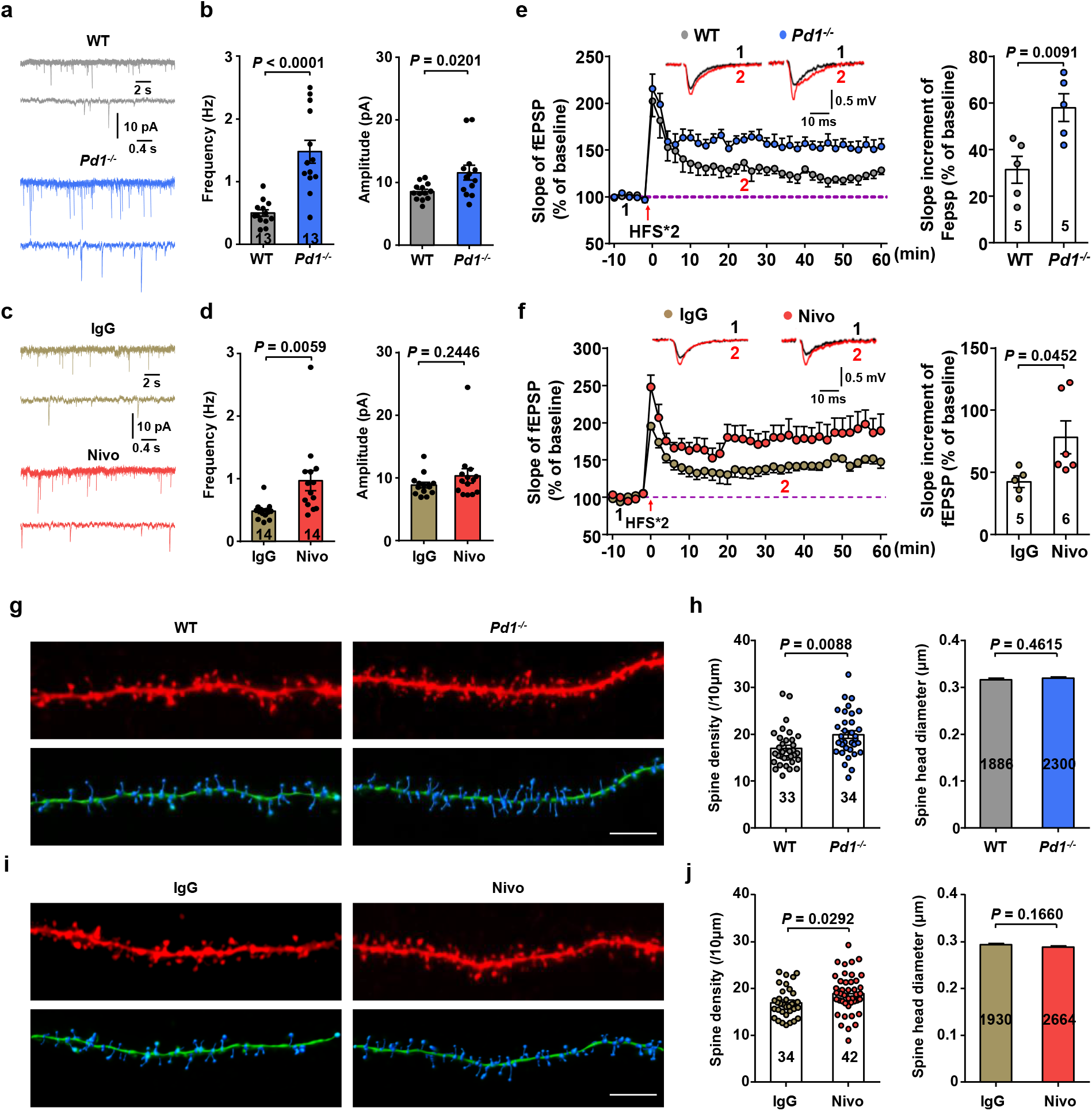
Enhanced synaptic plasticity and increased dendritic spine density in hippocampal CA1 neurons with *Pd1*^−/−^ deficiency or after PD-1 blockade with nivolumab. **(a)** Representative mEPSCs traces in CA1 neurons in brain slices of WT and *Pd1*^−/−^ mice. **(b)** Increased mEPSCs frequency and peak amplitude in CA1 neurons of *Pd1*^−/−^ mice. **(c)** Representative mEPSCs traces in CA1 neurons of IgG- and nivolumab (7 nM, 2 h) treated slices. **(d)** Increased mEPSCs frequency but not mEPSCs amplitude in CA1 neurons in nivolumab-treated slices. **(e)** Enhanced LTP in *Pd1*^−/−^ slices. Top: Black and red traces represent the baseline fEPSP and post-induction fEPSP traces, respectively. Right: summary data of the slope increment of fEPSP in WT and *Pd1*^−/−^ mice. **(f)** Enhanced LTP, as indicated by slope of fEPSP, in nivolumab (7 nM, 2 h) treated brain slices of WT mice. Top: Black traces (1) and red traces (2) represent the baseline fEPSP and post-induction fEPSP, respectively. Right: summary data of the slope increment of fEPSP in IgG- and nivolumab-treated slices. **(g, h)** *Pd1* deficiency increases spine density of CA1 neurons. **(g)** Representative confocal stacks and three-dimensional reconstruction images of the apical dendrites of CA1 neurons obtained from both WT and *Pd1*^−/−^ mice. Scale bar, 5 μm. **(h)** Increased spine density but unaltered spine head diameter in CA1 neurons of *Pd1*^−/−^ mice. **(i, j)** Nivolumab increases spine density of CA1 neurons. **(i)** Representative confocal stacks and three-dimensional reconstruction images of the apical dendrites of CA1 neurons obtained from both IgG- and nivolumab-treated mice. Scale bar, 5 μm. **(j)** Spine density but not spine head diameter is increased in CA1 neurons in nivolumab-treated mice. Two-tailed unpaired t test for **b, d, e, f, h, j.** Data were expressed as mean ± s.e.m.

To rule out the contribution of surrounding cells to the electrical activities recorded in the CA1 neurons, we also examined neuronal excitability in dissociated hippocampal neurons in primary cultures from E17-E19 mice. Compared to WT neurons (7-8 days in vitro, DIV), *Pd1*-deficient primary neurons (DIV 7-8) had a moderate increase in RMP (*P*=0.0729, **Fig. 2h**) but fired more AP following a ramp of current injection (*P*=0.0002, **Fig. 2i**). Notably, 39% (7/18) of *Pd1*-deficient neurons exhibited spontaneous discharges, while only 12.5% (2/16) WT neurons exhibited spontaneous discharges (**Fig. 2j**). Thus, CA1 neuron-intrinsic PD-1 is sufficient to regulate neuronal excitability.

We next examined synaptic transmission and plasticity in CA1 neurons in brain slices with PD-1 deletion or blockade. We found marked increases in the frequency (*P*<0.0001) and amplitude (*P*<0.05) of miniature EPSCs (mEPSCs) in CA1 neurons of KO mice (**Fig. 3a, b**). PD-1 blockade with nivolumab (7 nM, 2 h) in WT slices also increased mEPSC frequency (*P*<0.05, **Fig. 3c, d**). As an important form of synaptic plasticity underlying learning and memory^11^, long-term potentiation (LTP) in CA1 neurons could be induced by high frequency electrical stimulation of Schaffer collaterals in brain slices (**Fig. 3e, f**). Interestingly, this form of LTP was greatly enhanced in *Pd1*-deficient (*P*<0.01, vs. WT slices, **Fig. 3e**) as well as in WT brain slices treated with nivolumab (7 nM, 2 h, *P*<0.05, vs. IgG control, **Fig. 3f**).

LTP is also associated with dendritic morphological changes^12^. Loss of PD-1 resulted in increased dendritic spine density in CA1 neurons of KO mice (*P*<0.01, **Fig. 3g, h**), without affecting neuronal survival in the hippocampus (**Supplementary Fig. 4**). In agreement with the result from KO mice, anti-PD1 treatment in WT slices with nivolumab (7 nM, 2 h) also increased spine density (*P*<0.05, **Fig. 3i, j**). The size of spines did not change after *Pd1* deletion or PD-1 blockade (**Fig. 3g-h**).

In summary, our findings have demonstrated a previously unrecognized role of neuronal PD-1 in regulating learning and memory. Previous studies showed that systemic anti-PD-1 treatment enhanced memory in mouse AD models, as a results of recruitment of immune cells (e.g., macrophages) and subsequent clearance of amyloid-β plaques ^3, 4^. Contradictory results were also reported ^5, 6^. We previously demonstrated the presence of functional PD-1 in peripheral sensory neurons which regulates nociceptor excitability and pain^7^. In this study, we demonstrated that functional PD-1 is present in CNS hippocampal neurons and PD-1 serves as an endogenous inhibitor of learning and memory via a direct modulation of neuronal excitability and synaptic transmission. Thus, PD-1 is a neuronal inhibitor both in the PNS and CNS. Inhibition of PD-1 could enhance learning and memory in young mice and 10 month-old mice with age-related cognitive decline. Mechanistically, anti-PD-1 treatment potentiates learning and memory by enhancing neuronal excitability, synaptic transmission, and synaptic plasticity (LTP) and increasing dendritic spines in CA1 neurons. Our findings suggest that anti-PD-1 treatment, a magic bullet for immunotherapy, may also be used as a neurotherapy for enhancing learning and memory. Future studies are warranted to test whether anti-PD1 treatment enhances memory in AD and dementia patients via both immune- and neuro- therapies.

## METHODS

Methods are available in the online version of the paper.

*Note: Supplementary Information is also available in the online version of the paper.*

## ACKNOWLEDGEMENTS

This study is supported by NIH RO1 grant DE17794.

## AUTHOR CONTRIBUTIONS

J.Z. conducted behavioral, electrophysiological, histochemical and primary cell culture experiments, and prepared the figures. R.-R. J. supervised the project. R.-R.J. and J.Z. wrote the paper.

## COMPETING FINANICIAL INTERESTS

The authors declare no competing financial interests.

## Data availability statement

The original data are available upon the request.

---

**Supplementary Fig. 1.**
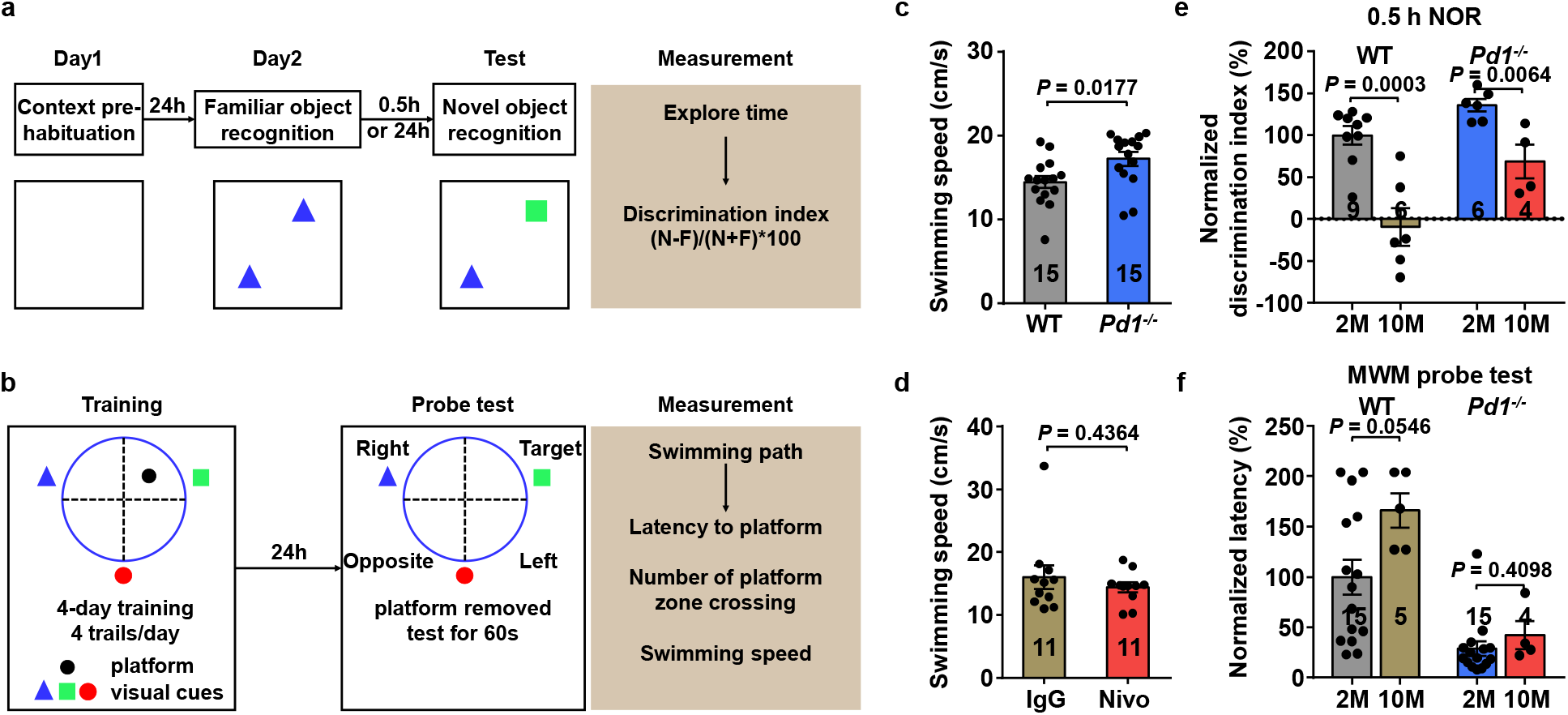
Novel object recognition test and Morris water maze test in mice. **(a)** Schematic of the novel object recognition procedures. **(b)** Schematic of the Morris water maze training and probe tests procedures. **(c, d)** Swimming speed of WT and *Pd1*^−/−^ mice **(c)** and in WT mice treated with IgG- and nivolumab **(d)** in Morris water maze probe tests. **(e)** Normalized discrimination index (% of young-WT mice) of young- and 10 month old-WT and *Pd1*^−/−^ mice in novel object recognition test. **(f)** Normalized latency to platform (% of young-WT mice) of young- and 10 month old -WT and *Pd1*^−/−^ mice in Morris water maze probe test. Two-tailed unpaired t-test for **c-f.** Data were expressed as mean ± s.e.m.

**Supplementary Fig. 2.**
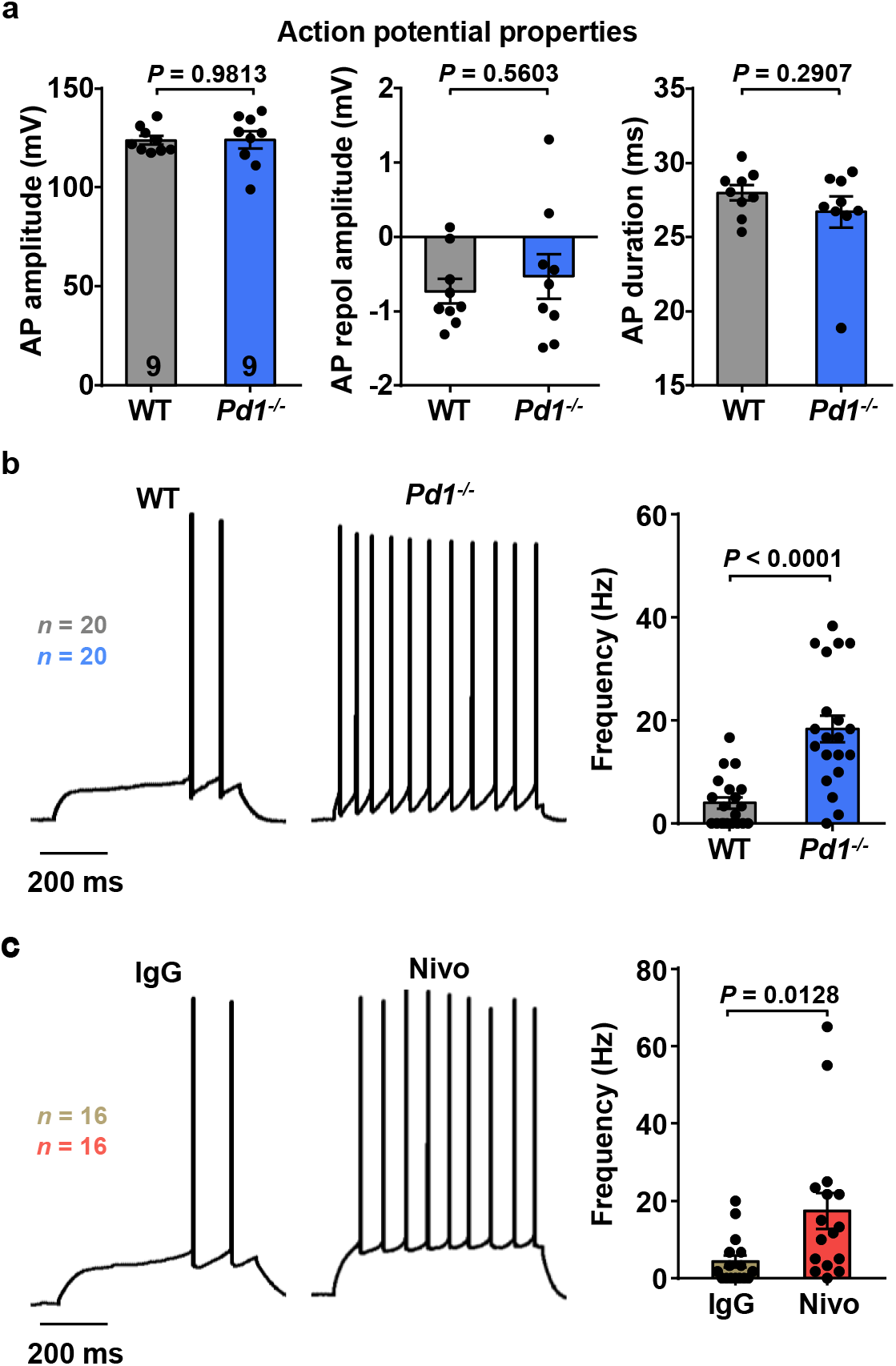
Evoked action potential in CA1 neurons of brain slices in mice. **(a)** Action potential properties (AP amplitude, AP repol amplitude, and AP duration) in CA1 neurons of WT and *Pd1*^−/−^ mice. **(b)** Representative AP traces and frequency in CA1 neurons of WT and *Pd1*^−/−^ mice, induced by 40 pA current. **(c)** Representative AP traces and frequency (induced by 40 pA current) in CA1 neurons of IgG- and nivolumab-treated slices of WT mice. Two-tailed unpaired t-test for **a-c**. Data were expressed as mean ± s.e.m.

**Supplementary Fig. 3.**
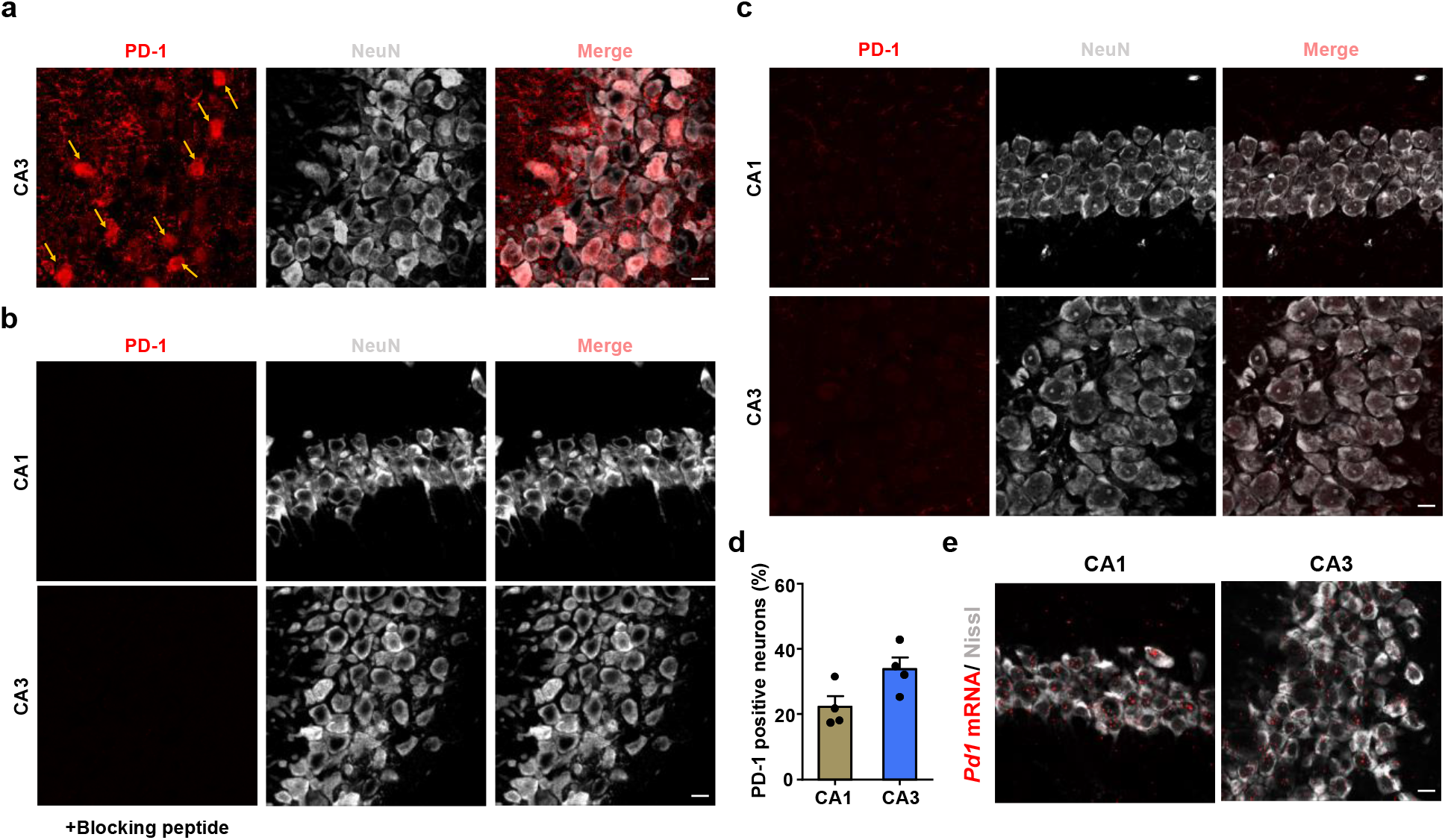
Expression of PD-1 and *Pd1* mRNA in mouse hippocampus neurons. **(a)** PD-1 is expressed by CA3 neurons. Arrows indicate PD-1 positive neurons. Scale bar, 10 μm. **(b)** PD-1 immunostaining signal is absent upon treatment with a blocking peptide in WT mice. Scale bar, 10 μm. **(c)** PD-1 immunostaining signal is absent in *Pd1*^−/−^ mice. Scale bar, 10 μm. **(d)** Percentage of PD-1 positive neurons in the CA1 and CA3 regions of the hippocampus. **(e)** *In situ* hybridization using RNAscope showing *Pd1* mRNA expression of CA1 and CA3 neurons of WT mice. Scale bar, 10 μm. Data were expressed as mean ± s.e.m.

**Supplementary Fig. 4:**
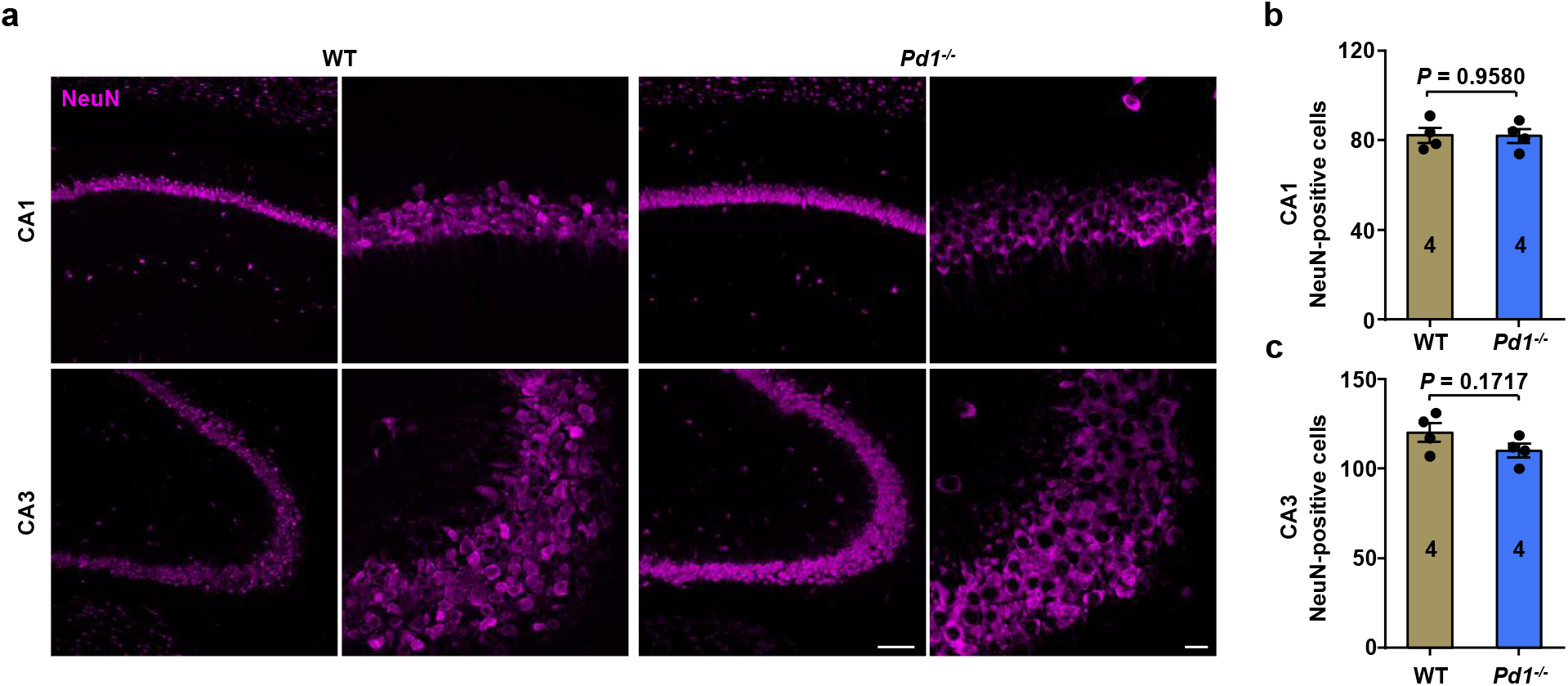
No neuronal loss in the hippocampal CA1 and CA3 regions of *Pd1*^−/−^ mice. **(a)** Confocal microscopy images of NeuN-positive neurons in CA1 and CA3 of WT and *Pd1*^−/−^ mice. Scale bar, 100 μm and 10 μm. **(b,c)** Quantification of the number of NeuN-positive neurons in CA1 **(b)** and CA3 **(c)** of WT and *Pd1*^−/−^ mice. Two-tailed unpaired t-test. Data were expressed as mean ± s.e.m.

## METHODS

### Animals

Adult mice (males, 2-3 months or 10 months) were used for behavioral test. Mice (5-8 weeks of both sexes) were used for electrophysiological studies. *Pd1* knockout mice with C57BL/6 background were purchased from the Jackson laboratory (Stock No: 021157) and maintained at Duke animal facility. All the mouse procedures were approved by the Institutional Animal Care & Use Committee of Duke University. All animals were housed under a 12-hour light/dark cycle with food and water available *ad libitum*. No statistical method was used to predetermine sample size. Sample sizes were estimated based on our previous studies for similar types of behavioral, biochemical, and electrophysiological analyses^13-15^. Animals were randomly assigned to each experimental group. Two to five mice were housed in each cage. All behavioral measurements were conducted in a blinded manner and during daytime (light cycle) normally starting at 9 AM. Animal experiments were conducted in accordance with the National Institutes of Health Guide for the Care and Use of Laboratory Animals. The number of animals and sample size of each experiment was also described in **Supplementary Table 1**.

### Reagents, stereotaxic surgery and drug delivery

Human IgG4 (Catalog: ab90286) were purchased from Abcam. Nivolumab (OPDIVO®), a humanized anti-PD-1 antibody, was purchased from Bristol-Myers Squibb. For stereotaxic surgery, mice were anesthetized with isoflurane (4% for induction and 2% thereafter for maintenance), and their heads were fixed in a stereotaxic apparatus (Narishige scientific instrument lab.). A guide cannula (62004, RWD Life Science) was stereotaxically implanted into the left lateral ventricles (AP: −0.3 mm; ML: +1.3 mm; V: −2.0 mm) based on the mouse brain atlas^16^. After four-day recovery, an infusion needle (62204, RWD Life Science) was inserted into lateral ventricles through the guide cannula to a depth of 2.5 mm for drug injection. Following the completion of the infusion, the needle was left for an additional 2 min to limit reflux.

### Novel object recognition test

On the first day, mice were placed in a 30 × 30 × 30 cm square arena (TAP plastics) for habituation. On the second day, two identical objects were placed in two distinct corners of the arena. Then, the mice were returned to its home cage for a 0.5-h or 24-h retention interval, and one of two identical objects was replaced by a new object for novel object recognition test^17^. After the retention interval, animals were placed back into the testing arena for 5-min exploration. A valid exploration was defined as mouse touching an object with nose or focusing attention toward the object at a distance of <1 cm. Turning around, climbing, and siting on the object were all considered to be invalid. A discrimination index was used to evaluate the scores of exploration time for the familiar or novel object (N - F/N + F) ×100%. Animal behaviors were video-recorded and analyzed by an experimenter who was blind to the testing conditions. Data were excluded if the total exploration time is less than 10 s.

### Morris water maze test

The Morris water maze test was conducted in a circular pool (diameter 120 cm) filled with 40 cm-height water maintained at room temperature (23 ± 1°C). The water was made opaque with white milk powder. The pool is located in an experimental room with many visual cues and divided into four equal quadrants. A circular platform, 10 cm in diameter, is placed in the middle of one fixed quadrant of the pool, just 1 cm underneath the water surface. During the training process, mice were trained for four trials each day for four consecutive days. In the probe test (day 5), the platform was removed and all mice were given one probe trial for 60 s searching^18^. The swimming speed, latency to platform zone and number of platform zone crossing were automatically recorded by ANY-maze (Stoelting Co.).

### Brain hippocampal slices preparation

Five to eight weeks old mice were anesthetized and then decapitated. The brain was quickly transferred to ice-cold artificial sucrose-based cerebrospinal fluid (ACSF) containing (in mM): Sucrose 75, NaCl 87, KCl 2.5, NaH_2_PO_4_ 1.25, CaCl_2_ 0.5, MgCl_2_ 7, NaHCO_3_ 26 and glucose 25. Hippocampal slices (200-300 μm) were cut using a vibratome (Leica VT1200S). Subsequently, slices were transferred to normal ACSF solution (in mM): NaCl 124, KCl 3, NaH_2_PO_4_ 1.25, CaCl_2_ 2, MgSO_4_ 1, NaHCO_3_ 26, glucose 10 at 34°C. All extracellular solutions were constantly carbogenated (95% O_2_, 5% CO_2_). Slices were kept in ACSF for at least 1 h before recording.

### Primary neuronal cultures of hippocampal neurons

Hippocampal neuron primary cultures were prepared from embryonic day 17-19 (E17-19) WT and *Pd1* knockout mice. Embryos were removed from maternal mice anesthetized with isoflurane and euthanized by decapitation. Hippocampi were dissected and placed in Ca^2+^- and Mg^2+^-free Hank’s balanced salt solution (HBSS, GIBCO) and digested at 37 °C in humidified O_2_ incubator for 30 min with collagenase type II (Worthington, 285 units/mg, 12 mg/ml final concentration) and dispase II (Roche, 1 unit/mg, 20 mg/ml) in HBSS (pH 7.3). Digestion was stopped by fetal bovine serum-added DMEM (GIBCO). Hippocampi were mechanically dissociated using fire-polished pipettes, filtered through a 100-μm nylon mesh and centrifuged (1,000 g for 5 min). The pellet was resuspended and plated on poly-D-Lysine/Laminin coated glass coverslips (CORNING), and cells were plated in DMEM containing 5% fetal bovine serum, 1% penicillin and streptomycin. After 5-6 h, primary cultures were switched to Neurobasal Plus medium containing 2% B27 supplement, 1% GlutaMAX-I, 1% penicillin and streptomycin (GIBCO). Three days after plating, cytosine arabinoside was added to a final concentration of 10 μM to curb glial proliferation. Whole-cell recoding and immunostaining experiments were conducted on DIV 7-8.

### Whole-cell recording of hippocampal slices

Whole-cell recordings were performed at 34 ± 1 °C with the help of an automatic temperature controller (Warner instruments) with an EPC10 amplifier (HEKA). Data were low-pass-filtered at 2 KHz, sampled at 10 KHz. Patch pipettes were filled with a solution containing the following (in mM): K-gluconate 135, KCl 5, CaCl_2_ 0.5, MgCl_2_ 2, HEPE 5, EGTA 5, MgATP 5 (pH 7.3, 290–300 mOsm/L). When filled with the pipette solution, the resistance of the pipettes was 4-8 MΩ. RMPs, spontaneous spikes and action potentials were recorded in current-clamp mode. The action potentials were evoked by current injection steps. Data were analyzed by Patchmaster (HEKA). For mEPSCs recording, neuron was hold at −70 mV in the presence of 0.5 μM TTX and 50 μM picrotoxin to block Na^+^ currents and GABA_A_ receptors. The mEPSCs were detected and analyzed using Mini Analysis (Synaptosoft Inc.). All drugs and regents were purchased from Sigma or Tocris.

### Extracellular field recordings of LTP in brain slices

To record extracellular field excitatory postsynaptic potentials (fEPSP), a bipolar tungsten stimulating electrode (FHC) was placed along the Schaffer collateral fibers to deliver test and conditioning stimuli, and a glass recording electrode (4–8 MΩ, filled with ACSF) was placed in the stratum radiatum of the CA1 region, 150–200 μm away from the stimulating electrode. The intensity of the stimulation was adjusted to produce an fEPSP with an amplitude of 30–40% of the maximum response. Test stimulation was delivered once per 30 s. Once a stable test response was attained, fEPSP baselines were recorded every 30 s for 10 min. Then, the LTP was induced by two HFS (100 Hz for 1 s, 30 s interval). After induction, fEPSP was recorded for another 60 min. The slope of fEPSP was analyzed by Patchmaster (HEKA).

### Visualization of dendrites in hippocampal neurons and morphological analysis

To visualize the dendritic spines of CA1 neurons, Neurobiotin (0.2%, Vector Laboratories) was dissolved in the intracellular pipette solution. After 30 min of whole-cell patch recording, the slices were fixed with 4% PFA in PBS and then processed using Alexa Fluor 594 streptavidin (1:500, Life Technologies, Catalog:S32356) for visualization. Neuronal dendrites and dendritic spines were imaged by ZEISS LSM 880 with Airyscan. Imaris software (v.9.3.0, Bitplane) was used to reconstruct dendritic spines and analyze the spine density and head diameter. The spine density was analyzed as previously reported (PMID: 31332372, PMID: 30685224, PMID: 29429933, PMID: 20037574, PMID: 24912492).

### Immunohistochemistry in mouse brains

After appropriate survival times, mice were deeply anesthetized with isoflurane and perfused with PBS, followed by 4% paraformaldehyde (PFA). After the perfusion, brain was removed and post-fixed in 4% PFA overnight and then dehydrated in 30% sucrose for 48 h. Brain tissue sections (14 μm) and free-floating brain sections (30 μm) were cut in a cryostat. The sections were blocked with 5% donkey serum for 1 h at room temperature and then incubated overnight at 4°C with the following primary antibodies: anti-PD-1 antibody (rabbit, 1:300, Sigma, Catalog: PRS4065) and anti-NeuN antibody (mouse, 1:1000, Millipore, Catalog: MAB377). After washing, the sections were incubated with cyanine 3(Cy3)- or Cy5-conjugated secondary antibodies (1:400; Jackson ImmunoResearch) at 4°C overnight. DAPI (1:1000, Thermo Scientific, Catalog: 62248) was used to stain cell nuclei. The stained sections were examined with a Leica SP5 inverted confocal microscope. To confirm the specificity of PD-1 antibody, blocking experiments were conducted in brain sections using a mixture of anti-PD-1 antibody (1:300) and immunizing blocking peptide (1:300, Sigma, Catalog: SBP4065), based on a protocol recommended for blocking with immunizing peptide (www.abcam.com/technical). Specificity of PD-1 antibody was also tested in brain sections of *Pd1* knockout mice. To determine if there is neuronal loss in *Pd1*^−/−^ mice, NeuN-positive neurons were quantified in CA1 and CA3 by Image J. Two sections were analyzed in each mice and 4 mice per group were analyzed.

### Immunocytochemistry in mouse primary hippocampal neurons

For immune staining in primary hippocampal neuron cultures (DIV 7-8) prepared from E17-19 WT and *Pd1*^−/−^ embryos, neurons grown on coverslips were fixed in 4% PFA for 30 min. After washing, the coverslips with neurons were blocked with 5% donkey serum for 1 h at room temperature and then incubated overnight at 4°C with the following primary antibodies: anti-PD-1 antibody (rabbit, 1:300, Sigma, Catalog: PRS4065) and anti-NeuN antibody (mouse, 1:1000, Millipore, Catalog: MAB377). Then, the sections were incubated with cyanine 3(Cy3)-and Cy5-conjugated secondary antibodies (1:400; Jackson ImmunoResearch) at 4°C overnight. DAPI (1:1000, Thermo Scientific, Catalog: 62248) was used to stain cell nuclei. The stained coverslips were examined with ZEISS LSM 880 inverted confocal microscope.

### In situ hybridization

We used probes directed against mouse *Pdcd1* (NM_008798) designed by Advanced Cell Diagnostics and the RNAscope multiplex fluorescent assay according to the manufacturer’s instructions. Pre-hybridization, hybridization and washing were performed according to standard methods^13^. Nissl staining (1:200, Thermo Fisher Scientific, Catalog: N21483) was used to stain neurons in tissue sections.

### Statistical analyses

All the data were expressed as mean ± s.e.m. The sample size for each experiment was based on our previous studies on such experiment^13, 15^. Statistical analyses were completed with Prism GraphPad 8.0. Biochemical, electrophysiology and behavioral data were analyzed using two-tailed unpaired t-test (two groups) or Two-Way ANOVA followed by Bonferroni post-hoc test. The criterion for statistical significance was *P* < 0.05.

### Data availability

The data that support the findings of this study are available from the corresponding author upon request.

